# Development of an Antiseizure Drug Screening Platform for Dravet Syndrome at the NINDS contract site for the Epilepsy Therapy Screening Program

**DOI:** 10.1101/2020.12.01.406470

**Authors:** Chelsea D. Pernici, Jeffrey A. Mensah, Elizabeth J. Dahle, Kristina J. Johnson, Laura Handy, Lauren Buxton, Misty D. Smith, Peter J. West, Cameron S. Metcalf, Karen S. Wilcox

## Abstract

**Objective:** Dravet syndrome (DS) is a rare, but catastrophic genetic epilepsy, with 80% of patients with carrying a mutation in the SCN1A gene. Currently, no anti-seizure drug (ASD) exists that adequately controls seizures. Patients with DS often present clinically with a febrile seizure and generalized tonic-clonic seizures that continue throughout life. To facilitate the development of ASDs for DS, the contract site of the NINDS Epilepsy Therapy Screening Program (ETSP) has evaluated a mouse model of DS using the conditional knock-in *Scn1a*^*A1783V/WT*^ mouse.

**Methods:** Survival rates and temperature thresholds for *Scn1a*^*A1783V/WT*^ were determined. Prototype ASDs were administered via intraperitoneal injections at the time-to-peak effect, which was previously determined, prior to the induction of hyperthermia-induced seizures. Protection was determined if ASDs significantly increased the temperature at which *Scn1a*^*A1783V/WT*^ mice seized.

**Results:** Approximately 50% of *Scn1a*^*A1783V/WT*^ survive to adulthood and all have hyperthermia-induce seizures. The results suggest that hyperthermia-induced seizures in this model of DS are highly refractory to a battery of ASDs. Exceptions were clobazam, tiagabine, and the combination of clobazam and valproic acid with add-on stiripentol, which elevated seizure thresholds

**Significance:** Overall, the data demonstrate the proposed model for DS is suitable for screening novel compounds for the ability to block hyperthermia-induced seizures and heterozygous mice can be evaluated repeatedly over the course of several weeks, allowing for higher throughput screening.

**Key Points:**

- *Scn1a*^*A1783V/WT*^ mice have a 50% survival rate and all have hyperthermia-induced seizures.
- Common DS treatments such as CLB and combinatorial therapy of CLB, VPA, and STP increase temperature thresholds in *Scn1a*^*A1783V/WT*^ mice.
- Sodium channel blockers, such as CBZ and LTG, decrease temperature thresholds of *Scn1a*^*A1783V/WT*^ mice as predicted.
- S*cn1a*^*A1783V/WT*^ mice are highly pharmacoresitant to common ASDs
- The *Scn1a*^*A1783V/WT*^ may be a useful preclinical drug screening platform for the treatment of DS.

## Introduction

Dravet Syndrome (DS) is a rare, catastrophic form of genetic epilepsy manifesting in the first year of life of a seemingly normal infant ^1^. In 80% of cases, the disease is caused by a mutation in the *SCN1A* gene, which encodes for the voltage-gated sodium channel Nav1.1^2^. There are over 900 distinct SCN1A mutations, with a large percent resulting from missense or frameshift mutations, leading to a loss of function in the sodium channel^3^. In mouse models, the *Scn1a* mutation has shown reduce sodium currents and excitability in inhibitory interneurons^4–7^. As a result, this causes an imbalance between inhibition and excitation, leading to general hyperexcitability. Seizures in humans are usually first induced by a fever or other increases in core temperature^2^. Spontaneous seizures will begin occurring within weeks of the initial seizure, becoming progressively worse and occurring more frequently^8^. Besides high seizure burden, DS also negatively impacts development and behavior and, in some cases, results in sudden unexpected death in epilepsy (SUDEP). DS is highly pharmacoresistant, with the primary treatment goal to decrease the seizure frequency and prevent status epilepticus^9^. Current treatment options fail to adequately address both seizures and the comorbidities associated with DS in many patients, and therefore discovering effective anti-seizure drugs (ASDs) is imperative. Typical first-line treatments for DS include valproate and clobazam, while stiripentol and topiramate are secondary lines of treatment, usually used congruently with valproate and clobazam^10–12^. Both cannabidiol and fenfluramine are promising new therapeutics in DS patients, with cannabidiol reducing motor seizures by ~40%^13, 14^ and fenfluramine reducing seizures by ~50%^15–17^. Some commonly used ASDs for seizure control, such as carbamazepine and lamotrigine, exacerbate seizures in both children and mice with DS conferring mutations^8, 18^. These sodium channel blockers worsen seizures in patients due to reduced SCN1A function in inhibitory neurons, thereby promoting hyperexcitability^8^. While there have been recent advances in treatment options for patients with DS, full seizure freedom has yet to be achieved. Clinically relevant animal models with good face, construct, and predictive validity are critically needed to successfully screen and discover effective, novel therapies.

An important goal at the National Institutes of Neurological Disorders and Stroke (NINDs) contract site for the Epilepsy Therapy Screening Program (ETSP) is to have well-characterized, reliable, and translatable models to facilitate the development of novel therapies for the treatment of epilepsy, including genetic epilepsies such as DS. In order to develop a screening model for DS for potential adoption by the program, mice with a mutation in *Scn1a* were obtained from Jackson Laboratories. The mutation mini-cassette includes lox P sites and, in the presence of Cre recombinase, results in an amino-acid substitution from alanine to valine at position 1783 (A1783V), previously reported in patients with DS . This mutation in mice^19, 20^. This mutation in mice results in both hyperthermia-induced seizures, which mimics febrile seizures seen in a clinical setting, and spontaneous, recurrent seizures^21–23^. In the present study, we show the potential of using the hyperthermia-induced seizure phenotype to rapidly screen novel therapeutics for the treatment of DS. To characterize this model, a survival analysis and a temperature threshold at which heterozygous (*Scn1a*^*A1783V/WT*^) mice seize was determined. Following these experiments, a battery of ASDs for the treatment of epilepsy was administered to identify the pharmacological characteristics of this preclinical model. These data will serve as a metric for the identification of novel anti-seizure compounds.

## Methods

### Animals

All animal care and experimental procedures were approved by the Institutional Animal Care and Use Committee of the University of Utah. Animal experiments were conducted in a manner consistent with Animal Research: Reporting of In Vivo Experiments (ARRIVE) guidelines (https://www.nc3rs.org.uk/arrive-guidelines). Experimental animals were generated by breeding a floxed stop male *Scn1a*^A1783V^ (B6(Cg)-Scn1atm1.1Dsf/J, Jax #026133) with a Sox2-cre (B6.Cg-Edil3^Tg(Sox2-cre)1Amc^/J) female mouse to produce both heterozygous (*Scn1a*^*A1783V/WT*^) and wild-type offspring. Both female and male heterozygous and age-matched wild type littermates were used for experiments (4-6 weeks of age). Mice were group housed in a pathogen free facility under a 12-h light/12-h dark light cycle and had access to food and water ad-libitum, except during hyperthermia-induced seizure experiments, in which case, they were transferred to the experimental room approximately one hour before testing.

### Hyperthermia-induced seizures

To evaluate the temperature at which *Scn1a*^*A1783V/WT*^ mice seize, mice were placed under a heat lamp and core temperature was gradually raised in a plexiglass chamber until a generalized seizure was observed or the temperature reached 42.5°C^18, 24–26^. After the procedure, mice were transferred to a cool, granite block to rapidly lower the core temperature and then returned to their home cages. Body temperature was monitored using a neonate rectal probe (Braintree Scientific, Inc, Braintree, MA) coupled to a TCAT-2LV controller (Physitemp Instruments, Inc, Clifton, NJ). Mice acclimated in the chamber for 5 minutes prior to experiments. Male and female *Scn1a*^*A1783V/WT*^ randomly received drug or vehicle prior to the test and staff were blinded to the treatment. The temperature at which mice seized was recorded. In experiments where wildtype mice were used, staff was also blinded to genotype.

### Drug preparation and administration

Carbamazepine, clobazam, clonazepam, phenobarbital, phenytoin, stiripentol, and valproic acid were purchased from Sigma. Levetiracetam, lamotrigine, tiagabine, topiramate, fenfluramine, and rufindamide were obtained from TCI America (Portland, OR, U.S.A). Cannabidiol was purchased from Cayman Chemical (Ann Harbor, MI, U.S.A). Ezogabine and lacosamide were purchased from Axon Medchem (Reston, VA, U.S.A). For acute administration, all drugs were prepared as 0.5% methyl cellulose (Sigma; St. Louis, MO, U.S.A) suspensions. All drugs were administered in a volume of 0.01 mL/g. The dose (mg/kg) for each drug is listed in **Table 1**. All drug compounds were administered and tested based on their time-of-peak effect (TPE), which was previously determined in the maximal electroshock seizure (MES) model or in the 6 Hz test (data not shown). Intraperitoneal (i.p.) injections were administered at the TPE for each drug, which are listed in **Table 1**, prior to testing. For 5-day sub-chronic administration, compounds were administered at the same time each day and, on the last day, was administered at the TPE prior to hyperthermia-induced seizure testing. For the triple therapy study, clobazam and stiripentol were administered at 1.0 hr as a single injection and valproic acid was administered 0.25 hr in a separate injection prior to testing. In all testing, experimenters were blinded to treatment group and mice randomly assigned to treatment group.

**Table 1.** List of prototype drugs evaluated in *Scn1a*^A1783V/WT^ mice following intraperitoneal administration.

| Drug | Dosing Strategy | Dose (mg/kg) | TPE (hr) |
| --- | --- | --- | --- |
| **Clobazam (CLB)** | Single | 2.5, 5, 10 | 1.0 |
| **Carbamazepine (CBZ)** | Single | 40 | 0.25 |
| **Cannabidiol (CBD)** | Single | 100, 200 | 1.0 |
| **Clonazepam (CLN)** | Single | 0.5 | 0.5 |
| **Ezogabine (EZG)** | Single | 20 | 1.0 |
| **Fenfluramine (FFA)** | Single | 10, 25 | 0.5 |
| **Lacosamide (LCM)** | Single | 15 | 0.5 |
| **Lamotrigine (LTG)** | Single, sub-chronic | 10, 20 | 1.0 |
| **Levetiracetam (LEV)** | Single | 100 | 1.0 |
| **Lorcaserin** | Single | 10 | 0.5 |
| **Phenobarbital (PHB)** | Single | 15 | 0.5 |
| **Retigabine** | Single | 20 | 1.0 |
| **Rufinamide** | Single | 32 | 0.5 |
| **Stiripentol (STP)** | Single | 100 | 1.0 |
| **Tiagabine (TGB)** | Single | 1.3, 3, 5 | 1.0 |
| **Topiramate (TPM)** | Single | 300 | 1.0 |
| **Valproic Acid (VPA)** | Single | 300 | 0.25 |
| **CLB + VPA** | Single | 5, 150 | 1.0, 0.25 |
| **CLB + VPA** | Single | 5, 75 | 1.0, 0.25 |
| **CLB + VPA + STP** | Single | 5, 75, 100 | 1.0, 0.25, 1.0 |
| **CLB + VPA + STP** | Single | 5, 75, 30 | 1.0, 0.25, 1.0 |

### Cross-over studies

For cross-over studies, an initial cohort of 4-week-old *Scn1a*^*A1783V/WT*^ mice were randomly assigned into 2-3 treatment groups, including one vehicle group. The following week, mice were crossed over to another treatment group. This process was repeated for a maximum of four weeks. At the completion of testing, each mouse had received all treatments.

### Statistical Analysis

Statistical comparisons were conducted using a Log-rank (Mantel-Cox) test. Significance was defined as a p-value < 0.05. All analysis was conducted with GraphPad Prism 8.0. Sample sizes and statistical results are reported in figure legends. Data is presented as mean ± SD.

## Results

### Female *Scn1a*^*A1783V/WT*^ mice exhibit greater mortality than male mice during development

The *Scn1a*^*tm1.1Dsf*^ mouse has a conditional knock-in mutation that is expressed when exposed to cre recombinase. For all experiments, *Scn1a*^*tm1.Dsf*^ male mice were bred to female *Sox2-cre* mice. As a result of cre expression in oocytes, the offspring are either heterozygous for the *Scn1a* mutation (*Scn1a*^*A1783V/WT*^) or wild type (WT). Both the heterozygous and wild-type offspring were assessed for survival and compared. Over 60 days, approximately 50% of the *Scn1a*^*A1784V/WT*^ mice survived, while no deaths were observed in wild type littermates (**Figure 1A**). When survival was evaluated based on sex, male *Scn1a*^*A1783V/WT*^ mice had a significantly higher survival rate (60.0%) than female mice (32.0%) by adulthood (**Figure 1B)**.

**Figure 1.**
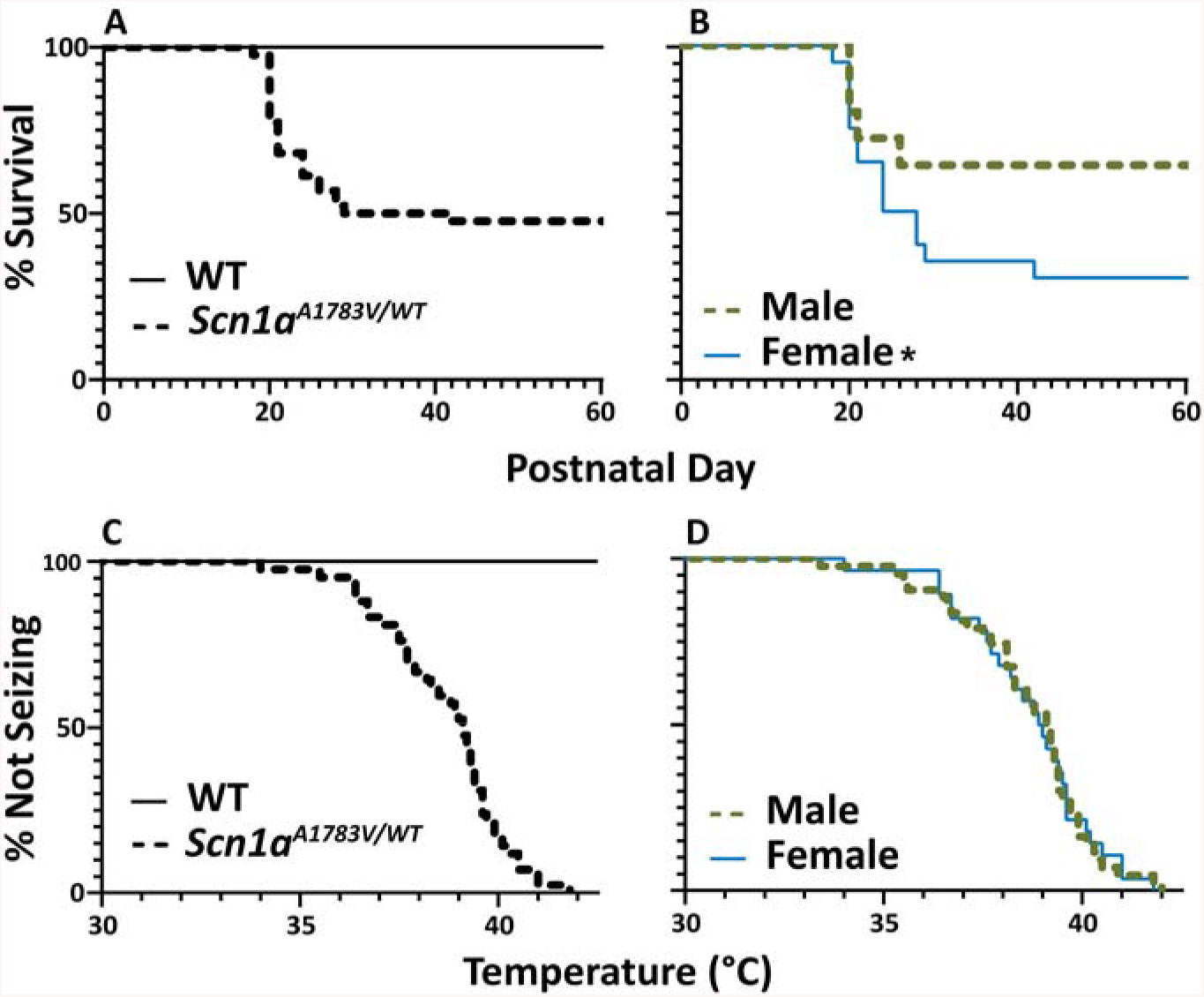
Approximately 50% of *Scn1a*^*A1783V/WT*^ mice survive to adulthood and female *Scn1a*^*A1783V/WT*^ mice exhibit greater mortality than male mice during development. **A)** By postnatal day 60, 51.1% of the observed *Scn1a*^*A1783V/WT*^ mice die, with the first death occurring at or around the age at which mice are weaned (P21). None of the wild type offspring died during the 60-day observation period (*Scn1a*^*A1783V/WT*^: n = 45, WT: n=30). **B)** When survival was evaluated based on sex, males had a significantly lower death rate (32.0%) than females (60.0%) by Day 60 (Males: n=25, Females: n = 20, p = 0.0397, Log-rank (Mantel-Cox)). **C)** All *Scn1a*^*A1783V/WT*^ mice had hyperthermia-induced seizures while age-matched wild type littermates had no evidence of seizure activity (*Scn1a*^*A1783V/WT*^: n = 71, WT: n = 16). **D)** There was no significant difference between the temperatures at which male and female *Scn1a*^*A1783V/WT*^ mice seized (Males: n =43, Females: n=28, p = 0.959, Log-rank (Mantel-Cox)).

### Male and female *Scn1a*^*A1783V/WT*^ mice have similar temperature thresholds for hyperthermia-induced seizures

The temperature threshold for which both male and female *Scn1a*^*A1783V/WT*^ mice seize was determined by subjecting mice to a steady increase in core temperature. Testing was discontinued when core temperature reached 42.5°C^18, 25, 26^. If no seizure occurred by 42.5°C, the mouse was considered seizure-free. All evaluated *Scn1a*^*A1783V/WT*^ mice had hyperthermia-induced seizures, with an average temperature threshold of 38.5 ± 1.9 °C. There was no evidence of seizure activity when the core temperature of wild type littermates was raised to 42.5°C (**Figure 1C**). To further evaluate if sex-dependent temperature thresholds exist, data was evaluated by sex. There was no significant difference between temperature thresholds for male (38.6 ± 1.8 °C) and female (38.7 ± 1.7 °C) *Scn1a*^*A1783V/WT*^ mice (**Figure 1D**).

### *Scn1a*^*A1783V/WT*^ mice can be used for several weeks of drug screening

To determine if *Scn1a*^*A1783V/WT*^ mice continue to have hyperthermia-induced seizures as they age and if mice could be used over the course of several weeks for drug screening of multiple therapeutics, a crossover testing scheme was utilized. Mice were randomly divided into three initial treatment groups (vehicle, CLB (10 mg/kg), and CBZ (40 mg/kg)). At each week, CLB, a drug which reduces seizure frequency in patients with DS^10^, significantly increased the temperature at which *Scn1a*^*A1783V/WT*^ mice had hyperthermia-induced seizures as compared to both vehicle and CBZ treated mice. On the other hand, CBZ, a drug which worsens seizures in patients with DS^10^, significantly decreased the temperature threshold at which *Scn1a*^*A1783V/WT*^ mice seized each week (**Figure 2**). Repeat treatment and repeat hyperthermia-induced seizures did not significantly affect the temperature at which *Scn1a*^*A1783V/WT*^ mice seized (**Supporting Figure 1**) in each treatment group.

**Figure 2.**
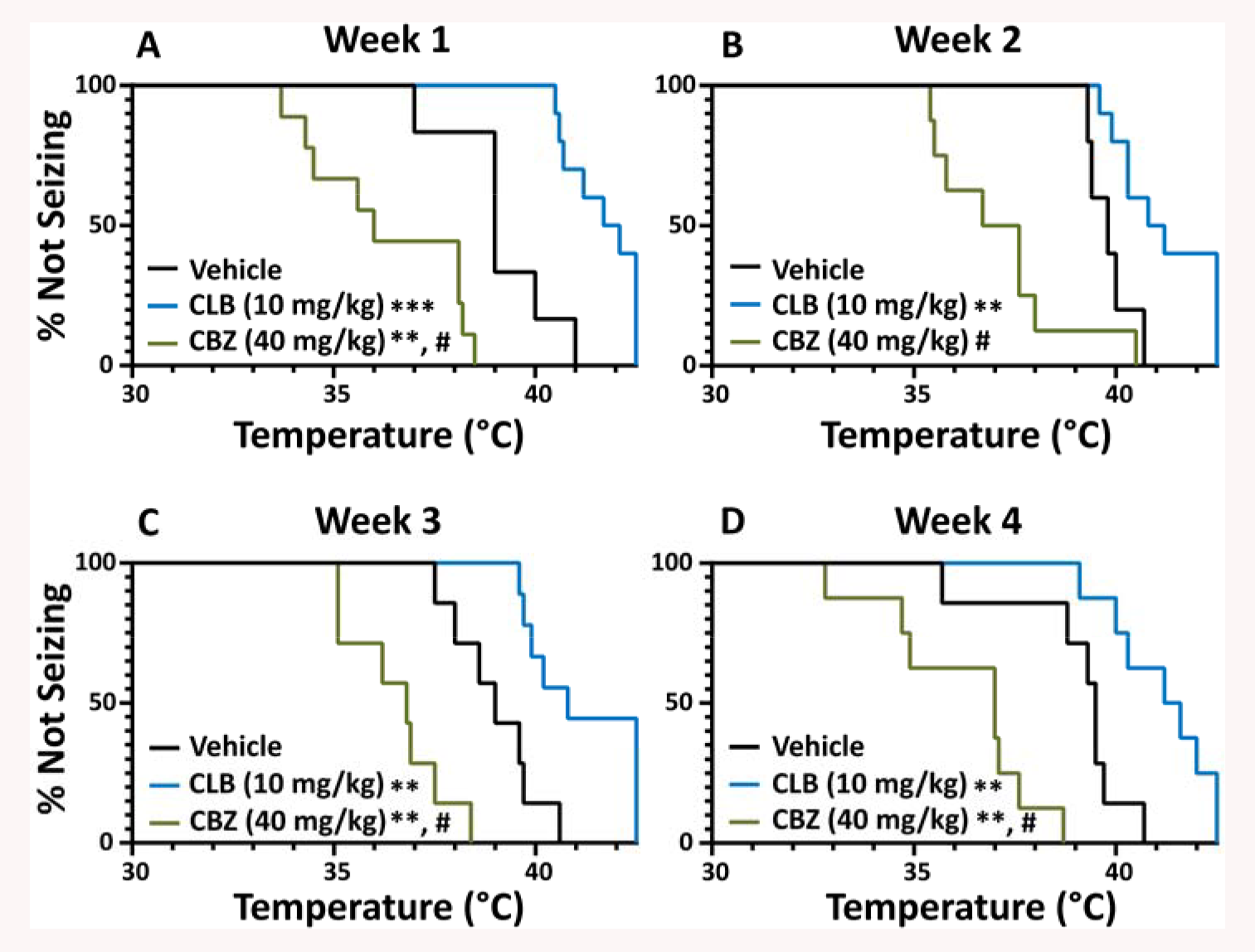
Repeat testing in a weekly crossover design does not change pharmacological outcome of seizures in *Scn1a*^*A1783V/WT*^ mice. *Scn1a*^*A1783V/WT*^ mice were randomly divided into three initial treatment groups (vehicle, CLB, and CBZ). Each week, mice were given a treatment not previously administered. At each week, CLB significantly increased the temperature threshold for which mice had hyperthermia induced seizures when compared to vehicle treated mice (Week 1 p = 0.0005, Week 2 p = 0.0076, Week 3 p = 0.0027, Week 4 p = 0.0048, Log-rank (Mantel-Cox)). Alternatively, CBZ decreased the temperature threshold each week, with a significant reduction seen at Week 1, 3, and 4 (Week 1 p = 0.0022, Week 2 p = 0.051, Week 3 p = 0.0014, Week 4 p = 0.0053, vs. vehicle, Log-rank (Mantel-Cox)). At each week, CLB significantly increased the temperature threshold as compared to CBZ (Week 1-4, p <0.0001, Log-rank (Mantel-Cox)). Vehicle: Week 1: n = 6, Week 2: n = 4, Week 3: n = 7, Week 4: n =7; CLB: Week 1: n = 10, Week 2: n = 10, Week 3: n = 9, Week 4: n =8; CBZ: Week 1: n = 9, Week 2: n = 8, Week 3: n =7, Week 4: n =8; ** p < 0.01, *** p < 0.001 vs. vehicle, #p < 0.0001 vs. CLB.

### Evaluation of prototype ASDs against hyperthermia-induced seizures

Additional ASDs that have previously shown efficacy in the treatment of DS, or those which are known to worsen seizure outcomes were evaluated against hyperthermia-induced seizures to determine the predictive validity of the *Scn1a*^*A1783V/WT*^ mouse model. Of those drugs known to confer seizure protection in DS, both CBD and FFA were evaluated^13, 27, 28^. CBD (200 mg/kg) did not significantly increase the temperature at which hyperthermia-induced seizures occurred (**Table 2**). FFA was evaluated at two doses, 10 mg/kg and 25 mg/kg. Neither dose had a significant effect on temperature threshold in *Scn1a*^*A1783V/WT*^ mice (**Supporting Figure 2 A&B, Table 2**).

**Table 2.**
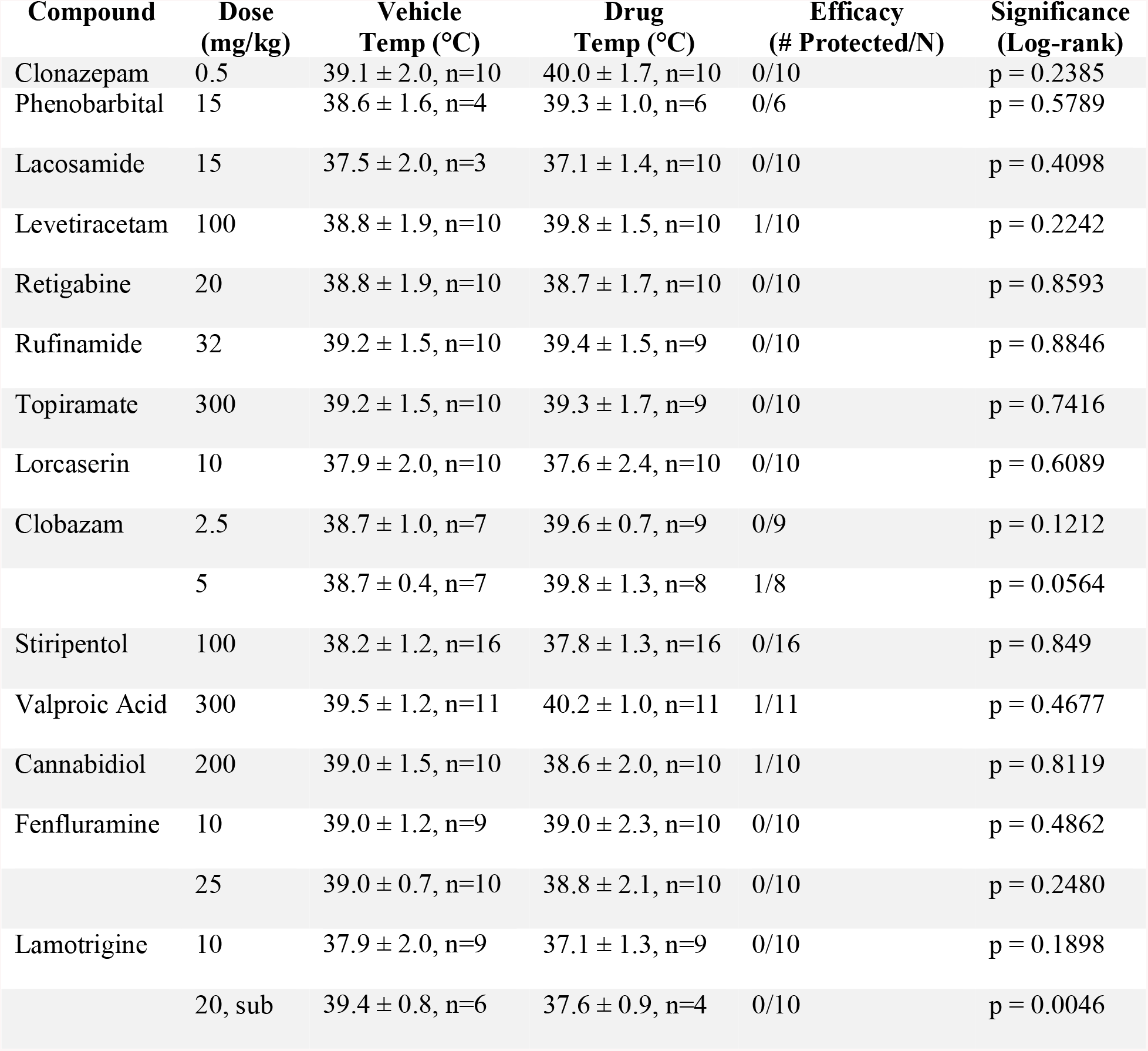
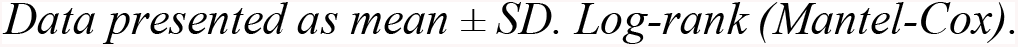
Anti-seizure drugs refractory to hyperthermia-induced seizures in *Scn1a*^*A1783V/WT*^ mice.

| **Compound** | **Dose (mg/kg)** | **Vehicle Temp (°C)** | **Drug Temp (°C)** | **Efficacy (# Protected/N)** | **Significance (Log-rank)** |
| --- | --- | --- | --- | --- | --- |
| Clonazepam | 0.5 | 39.1 ± 2.0, n=10 | 40.0 ± 1.7, n=10 | 0/10 | p = 0.2385 |
| Phenobarbital | 15 | 38.6 ± 1.6, n=4 | 39.3 ± 1.0, n=6 | 0/6 | p = 0.5789 |
| Lacosamide | 15 | 37.5 ± 2.0, n=3 | 37.1 ± 1.4, n=10 | 0/10 | p = 0.4098 |
| Levetiracetam | 100 | 38.8 ± 1.9, n=10 | 39.8 ± 1.5, n=10 | 1/10 | p = 0.2242 |
| Retigabine | 20 | 38.8 ± 1.9, n=10 | 38.7 ± 1.7, n=10 | 0/10 | p = 0.8593 |
| Rufinamide | 32 | 39.2 ± 1.5, n=10 | 39.4 ± 1.5, n=9 | 0/10 | p = 0.8846 |
| Topiramate | 300 | 39.2 ± 1.5, n=10 | 39.3 ± 1.7, n=9 | 0/10 | p = 0.7416 |
| Lorcaserin | 10 | 37.9 ± 2.0, n=10 | 37.6 ± 2.4, n=10 | 0/10 | p = 0.6089 |
| Clobazam | 2.5 | 38.7 ± 1.0, n=7 | 39.6 ± 0.7, n=9 | 0/9 | p = 0.1212 |
|  | 5 | 38.7 ± 0.4, n=7 | 39.8 ± 1.3, n=8 | 1/8 | p = 0.0564 |
| Stiripentol | 100 | 38.2 ± 1.2, n=16 | 37.8 ± 1.3, n=16 | 0/16 | p = 0.849 |
| Valproic Acid | 300 | 39.5 ± 1.2, n=11 | 40.2 ± 1.0, n=11 | 1/11 | p = 0.4677 |
| Cannabidiol | 200 | 39.0 ± 1.5, n=10 | 38.6 ± 2.0, n=10 | 1/10 | p = 0.8119 |
| Fenfluramine | 10 | 39.0 ± 1.2, n=9 | 39.0 ± 2.3, n=10 | 0/10 | p = 0.4862 |
|  | 25 | 39.0 ± 0.7, n=10 | 38.8 ± 2.1, n=10 | 0/10 | p = 0.2480 |
| Lamotrigine | 10 | 37.9 ± 2.0, n=9 | 37.1 ± 1.3, n=9 | 0/10 | p = 0.1898 |
|  | 20, sub | 39.4 ± 0.8, n=6 | 37.6 ± 0.9, n=4 | 0/10 | p = 0.0046 |

*Data presented as mean ± SD. Log-rank (Mantel-Cox).*

Sodium channel blockers such as CBZ and LTG can increase the frequency or severity of seizures in patients with DS^10^. 40 mg/kg CBZ significantly lowered the temperature at which hyperthermia-induced seizures occurred (**Figure 2A-D**). LTG was tested as a single 10 mg/kg dose and had no effect on the temperature threshold (**Supporting Figure 2C, Table 2**). However, when 20 mg/kg LTG was sub-chronically administered daily for 5 days and a hyperthermia-induced seizure was conducted on Day 5, it significantly lowered the temperature threshold as compared to vehicle treated mice. Additionally, during the 5-Day administration of LTG, 2 of the 6 mice in the treatment group died (**Supporting Figure 2D**).

### Except for tiagabine, hyperthermia-induced seizures in *Scn1a*^*A1783V/WT*^ mice are refractory to a battery of anti-seizure drugs

In addition to the compounds evaluated above, other prototype ASDs used in the treatment of epilepsy were evaluated. Of these additional compounds, only TGB was found to be effective in shifting the temperature threshold at which seizures occurred. TGB, which has shown to be protective against hyperthermia-induced seizures in a haploinsufficient mouse model of DS^29^ was evaluated at 1.3 mg/kg, 3 mg/kg, and 5 mg/kg. At all doses, TGB significantly increased the temperature at which mice seized as compared to vehicle (**Figure 3A-C**). At 1.3 mg/kg, 1 out of 10 treated mice was fully protected, at 3 mg/kg, 3 out of 9 treated mice were fully protected and at 5 mg/kg, 3 out of the 9 treated mice had complete protection against hyperthermia-induced seizures. None of the other ASDs listed in **Table 2** significantly affected the temperature threshold at which *Scn1a*^*A1783V/WT*^ mice had a hyperthermia-induced seizure (p > 0.05, Log-rank (Mantel-Cox)), suggesting this model of DS is highly refractive to a battery of ASDs.

**Figure 3.**
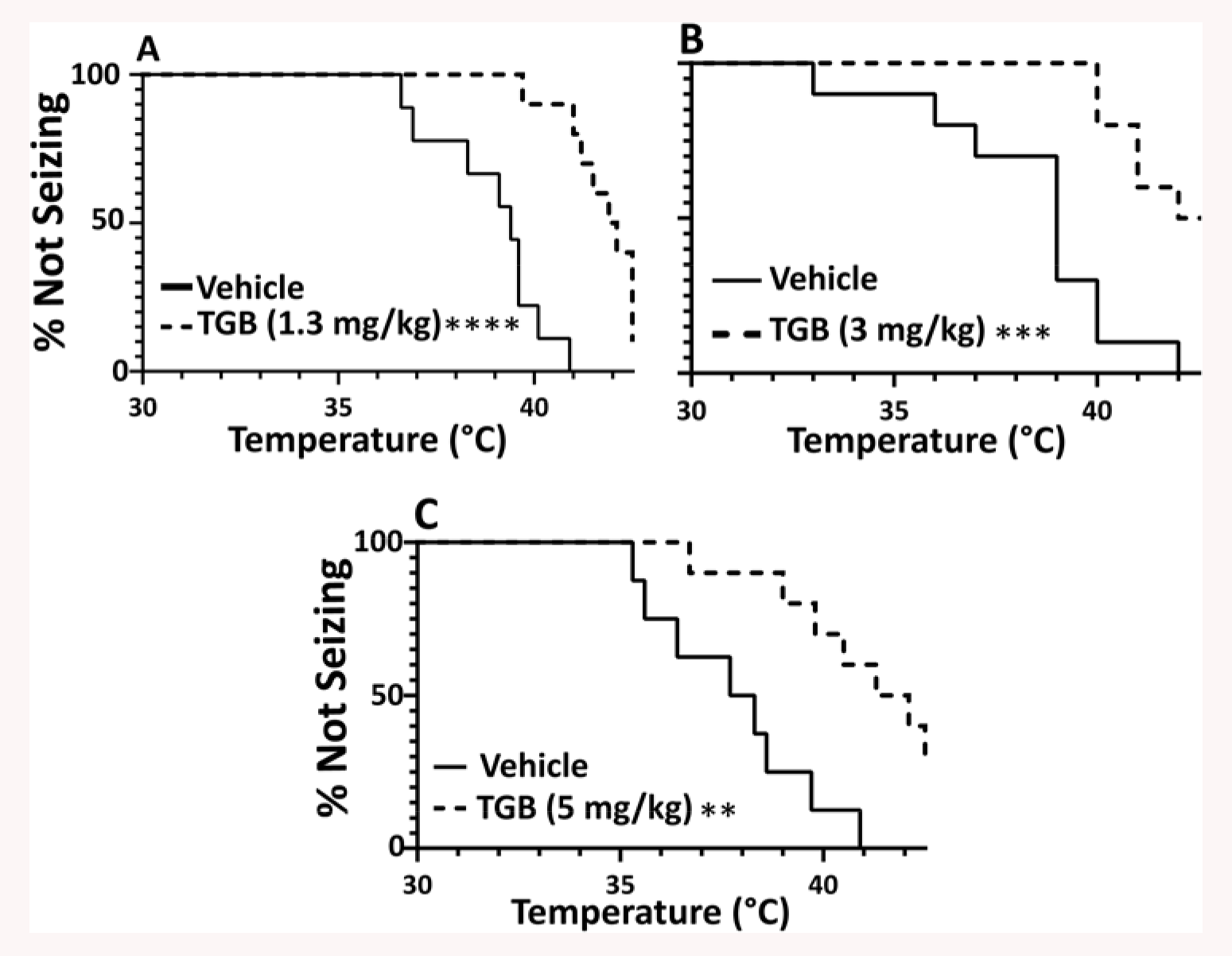
Evaluation of TGB. TGB was evaluated at three different doses to determine the temperature thresholds for hyperthermia-induced seizures. **A)** A dose of 1.3 mg/kg significantly increased the temperature threshold at which mice seized as compared to vehicle (p<0.0001), with 1/10 mice completely protected against hyperthermia-induced seizures. Vehicle: n=9; TGB (1.3 mg/kg): n=10 **B)** A dose of 3 mg/kg significantly increased the temperature at which mice had hyperthermia-induced seizures as compared to vehicle (p=0.0003), with 3/9 mice demonstrating complete protection. Vehicle: n=10; TGB (3 mg/kg): n=9 **C)** A dose of 5 mg/kg significantly increased the temperature at which mice had seizures (p=0.0014), with 3/10 showing complete protection. Vehicle: n=9; TGB (5 mg/kg): n=10 ** p < 0.01, *** p < 0.001, **** p < 0.0001; Log-rank (Mantel-Cox).

### Evaluation of add-on treatment against hyperthermia-induced seizures

First line treatment of DS includes either CLB or VPA^30^. When either fails as independent treatment, CLB and VPA are administered together^11^. Stand-alone treatment of either 5 mg/kg CLB or 300 mg/kg VPA did not offer significant protection against hyperthermia-induced seizures in *Scn1a*^*A1783V/WT*^ mice (**Table 2**). Therefore, as done in the clinic, CLB and VPA were administered congruently. 5 mg/kg CLB and 150 mg/kg VPA significantly raised the temperature threshold compared to vehicle treated mice (**Figure 4A**). If this combination fails in the clinic, a relevant second-line drug regimen combines CLB and VPA with STP as an add-on drug^11^. We tested a combination of 5 mg/kg CLB and 75 mg/kg which did not offer significant protection (**Figure 4B).** To mimic what is done in the clinic, 100 mg/kg STP was then “added on” to 5 mg/kg CLB and 75 mg/kg VPA treatment. The CLB (5 mg/kg) + VPA (75 mg/kg) + STP (100 mg/kg) treated mice had significantly higher temperature thresholds than CLB (5 mg/kg) + VPA (75 mg/kg) treated *Scn1a*^*A1783V/WT*^ mice (**Figure 4C**). Finally, we show that a lower dose of add-on STP does significantly raise temperature thresholds as compared to the combination of CLB (5 mg/kg) and VPA (75mg/kg) (**FIGURE 4D**). Thus, this add-on screening approach may prove useful when screening novel compounds for efficacy against hyperthermia induced seizures.

**Figure 4.**
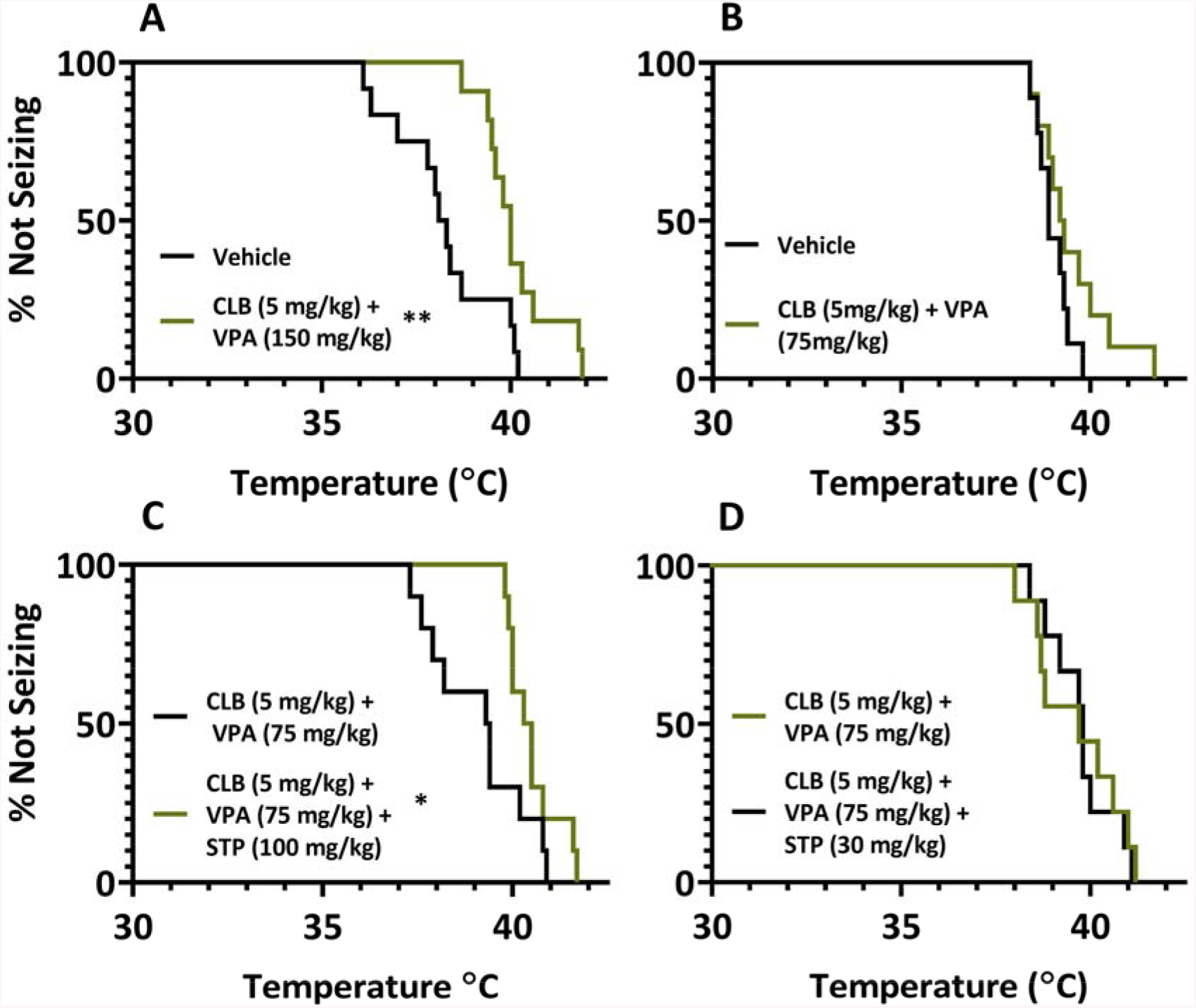
Add-on therapy for the treatment of Dravet Syndrome. **A)** CLB (5 mg/kg) and VPA (150 mg/kg) treatment significantly lowered the temperature threshold of hyperthermia-induced seizures as compared to vehicle treated mice (Vehicle: n= 12, CLB + VPA: n=11, p=0.0060). **B)** CLB (5 mg/kg) and VPA (75 mg/kg) did not significantly lower the temperature at which *Scn1a*^*A1783V/WT*^ mice had hyperthermia-induced seizures as compared to vehicle treated mice (Vehicle: n= 9, CLB+VPA: n=10, p=0.1511) **C)** The three drug combo of CLB (5 mg/kg), VPA (75 mg/kg) and STP (100 mg/kg) offered significant protection against hyperthermia-induced seizures as compared to CLB (5 mg/kg) + VPA (75 mg/kg) treated *Scn1a*^*A1783V/WT*^ mice (CLB+VPA+STP: n=10, CLB + VPA: n=10, p=0.0471). **D)** CLB (5 mg/kg) + VPA (75 mg/kg) + STP (30 mg/kg) did not significantly the temperature at which mice seized as compared to CLB (5 mg/kg) + VPA (75 mg/kg) treated mice (CLB + VPA + STP: n=9, CLB + VPA: n=9 p=0.7973) * p< 0.05, ** p < 0.01; Log-rank (Mantel-Cox).

## Discussion

Drug resistant epilepsies, like DS, critically need preclinical drug screening models to identify effective compounds. There are over 900 distinct mutations that result in this highly complex encephalopathy, further necessitating the availability of translatable models to discover effective therapies. Introducing new screening models to the Epilepsy Therapy Screening Program requires understanding both the face and predictive validity of proposed models. The *Scn1a*^*A1783V/WT*^ model has a survival rate of ~50% (**Figure 1A&B**), demonstrating the *Scn1a*^*A1783V/WT*^ mouse model has at least as good or better viability than other models, increasing the yield of experimental mice^4, 22, 31^. *Scn1a*^*A1783V/WT*^ mice maintain the hyperthermia-induced seizure phenotype (**Figure 1C&D**) as they age (**Figure 2**). While males have a significantly higher survival rate than females (**Figure 1B**) there was no difference in the temperatures at which they seized (**Figure 1D**). The breeding scheme used here allows for approximately half the offspring to be heterozygous for the mutation. This allows for faster breeding than in other models, especially as the breeding pairs, who do not express the floxed transgene, survive readily. The predictive validity of this model should be well characterized prior to screening novel compounds, as this provides a baseline metric from which to determine if novel compounds have superior efficacy over available compounds. Clinically approved ASDs for the treatment of DS, ASDs that are contradicted, as well as a battery of other ASDs, were screened to pharmacologically profile the *Scn1a*^*A1783V/WT*^ mouse model.

A first-line treatment for DS is CLB^10^. We found 10 mg/kg CLB significantly increased the temperature at which *Scn1a*^*A1783V/WT*^ mice seized (**Figure 2**). CLB has also been tested hyperthermia-induced seizures in the *F1.Scn1a*^*tm1Kea*^ mouse and 5 mg/kg CLB conferred protection^18^. In another study, single injections of CLB at 1 mg/kg and 10 mg/kg both significantly increased the temperature threshold of the *F1.Scn1a*^*tm1Kea*^ mouse^26^. In a mouse model with a truncation mutation, *Scn1a*^*R1407X*^, 6.62 mg/kg CLB also significantly increased the temperature at which mice had hyperthermia-induced seizures^32^. Overall, CLB has been effective against spontaneous seizures in patients^30^ and against hyperthermia-induced seizures in, along with the data obtained herein, three DS mouse models.

CBD, which was recently approved by the FDA as an adjunct to conventional ASDs, significantly reduces seizure frequency in patients^13, 28^. In the clinic, CBD is used as add-on therapy to currently prescribed treatment, with 66% of patients using it concomitantly with CLB^28^. In this present study, we tested CBD as a monotherapy and found 200 mg/kg CBD did not offer significant protection (**Table 2)**. While there was no protective effect in this model at the dose and time point tested, CBD monotherapy showed significant protection in two other DS mouse models. When 100 mg/kg CBD was administered to the F1.*Scn1a*^*tm1Kea*^ mouse it significantly increased the temperature at which mice seized^26^ and when administered to the *Scn1a*^+/−^ null mouse at both 100 mg/kg and 200 mg/kg, CBD significantly reduced seizure length and severity^33^. In the F1.*Scn1a*^*tm1Kea*^ mouse, the combination of 10 mg/kg CLB and 12 mg/kg CBD significantly increased the temperature threshold as compared to CBD monotherapy, suggesting the combination therapy may be more beneficial. Additionally, as not all people with DS have their seizures suppressed by CBD, not all mouse models of DS are protected from hyperthermia-induced seizures.

When treating DS, sodium channel blockers, such as CBZ and LTG, are avoided, as these can make seizure frequency and severity worse^9, 34^. In *Scn1a*^*A1783V/WT*^ mice, a single injection of 40 mg/kg CBZ significantly lowered the temperature at which mice seized (**Figure 2**). While a single dose of 10 mg/kg LTG had no effect on threshold temperatures (**Supporting Figure 2C, Table 2**), sub-chronic administration of 20 mg/kg LTG over the course of 5 days significantly lowered the temperatures at which mice seized and two mice died due to treatment (**Supporting Figure 2D, Table 2**). Both CBZ and LTG were tested in the F1.*Scn1a*^*tm1Kea*^ model, but a single dose of 20 mg/kg LTG significantly lowered temperature threshold, while 20 mg/kg CBZ had no effect. While our study did not include single administration of 20 mg/kg LTG, we presume it would also lower the temperature threshold, much like observed in the F1. *Scn1a*^*tm1Kea*^. The CBZ doses differed between the two studies, which could contribute to the difference seen in the temperature thresholds. However, given that our results align with what is seen in the clinical setting these compounds functioned as hypothesized in the *Scn1a*^*A1783V/WT*^ mouse model.

A notable effect in the present study was TGB conferred protection against hyperthermia-induced seizures in *Scn1a*^*A1783V/WT*^ (**Figure 3**). This is consistent with the pharmacological profile in a haploinsufficiency mouse model, as a high dose (10 mg/kg) significantly protected mice against generalized tonic-clonic seizures during hyperthermia testing^29^. While we saw significant protection in our model, with minimal toxic side effects, in patients, TGB is associated with behavioral side effects and reports of nonconvulsive absence and myoclonic seizures associated with TGB treatment^35^. Therefore, NICE guidelines recommend against using TGB for treatment of DS.

Other compounds used to treat some patients with DS, such as LEV and TPM, did not significantly confer protection in the *Scn1a*^*A1783V/WT*^ model (**Table 2**). However, in the F1.*Scn1a*^*tm1Kea*^ mouse model, 10 mg/kg LEV significantly increased temperature thresholds and 40 mg/kg TPM also showed no effect. LEV is only used when no other first-line treatment is successful^30^, with responder rates anywhere between 11%-64%^36, 37^. It’s not as surprising to discover this particular compound was effective in one mouse model and not the other, as there seems to be variability in the efficacy of this particular drug.

While CLB is commonly used in the treatment of DS, it’s responder rate is ~28%^30^ and add-on treatments are normally required to obtain adequate seizure control. These include the addition of drugs such as VPA and STP. 300 mg/kg VPA monotherapy did not offer significant protection as compared to vehicle treated mice (**Table 2).** Thus, we combined 5 mg/kg CLB and 150 mg/kg VPA and found it conferred significant protection (**Figure 4A**). We decreased the dose of VPA to 75 mg/kg, while maintaining CLB at a dose of 5 mg/kg and found it no longer significantly protected *Scn1a*^*A1783V/WT*^ mice (**Figure 4B**). When the combination of CLB and VPA fails in the clinic, STP is added to the treatment regimen^11^. We administered 5 mg/kg CLB, 75 mg/kg VPA, and 100 mg/kg STP and found this significantly offered protection as compared to 5 mg/kg CLB and 75 mg/kg VPA treatment (**Figure 4C**). We found this triple drug therapy to be effective and supports this pharmacological treatment regimen for DS. The combined use of CLB and VPA with an add-on investigational compound may be a useful approach towards novel therapy screening in this mouse model of DS.

Additional compounds were tested against hyperthermia-induced seizures in the *Scn1a*^*A1783V/WT*^ mouse model. Much like what is observed in the clinical setting with patients, this model is highly pharmacoresistant. However, as the majority of these experiments were all performed with acute injections, it might prove useful to determine the effect of these compounds when sub-chronically administered at doses sufficient to provide steady state therapeutic brain and plasma levels. Nevertheless, those experiments are technically challenging given the stress of multiple injections per day over the course of several days. Therefore, sub chronic dosing, wherein drugs can be delivered in food, would be a potential approach to addressing the limitations inherent in performing sub chronic dosing experiments with injections.

Another phenotype of DS is spontaneous, recurrent seizures. Drugs effective in patients, such as treatment with CBD (**Table 2**) and FFA (**Supporting Figure 2A&B, Table 2**), may be effective at reducing spontaneous seizure frequency in this mouse model or require chronic dosing in hyperthermia studies. In fact, when a patient presents in the clinic with a febrile seizure, they are typically administered rescue treatment, such as diazepam or midazolam. If patients are then diagnosed with DS, they are started on regimen of ASDs. Therefore, because a drug was not protective against hyperthermia-induced seizures does not necessarily mean it will not be effective against spontaneous seizures. Studies where conventional ASDs were screened in other mouse models of DS, variable results between hyperthermia-induced and spontaneous seizures were achieved. In the F1.*Scn1a*^*tm1Kea*^ mouse model, CLB was protective against hyperthermia-induced seizures, but had minimal effect against spontaneous seizures^18^. Surprisingly, in the F1.*Scn1a*^*tm1Kea*^, mouse, CBZ had no effect on either hyperthermia-induced or spontaneous seizures, while we saw a significant effect in lowering the temperature threshold of hyperthermia-induced seizures^18^. In another study using the F1*.Scn1a*^*tm1Kea*^ mouse, monotherapy CBD significantly increased the temperature threshold of mice, yet had no effect on spontaneous seizure frequency^26^. Therefore, it’s possible there is a pharmacological difference between hyperthermia-induced and spontaneous seizures, which may explain a lack of translation of some compounds to the clinic. In order to find effective therapies, novel therapeutics may also need to be screened against spontaneous seizures. Nonetheless, using hyperthermia-induced seizures is still an important screening platform as due to its highly pharmaco-resistant nature, we may be able to uncover extremely effective drugs.

Beyond the differences that may exist between hyperthermia-induced and spontaneous seizures at a pharmacological level, there are over 900 distinct mutations that result in DS^20^, making this form of epilepsy highly patient specific. Consequently, it is possible that mice will respond differently to ASDs when exhibiting different mutations. This could explain why some of the ASDs have shown to be effective against hyperthermia-induced seizures in the present model, which results from a missense mutation, and not in the F1.*Scn1a*^*tm1Kea*^ mouse, a model of haploinsufficiency. Therefore, we not only need translatable models for hyperthermia-induced and spontaneous seizures, but it would also be beneficial to have well characterized models for the different mutations.

An additional approach to screening novel compounds for DS is the use of zebrafish larvae. While zebrafish larvae do not recapitulate all the aspects of DS, such as cognitive disabilities, the *scn1a*^s552^ larvae exhibit unprovoked seizures^38^. This particular model has been validated with a number of ASDs and suppression of unprovoked seizure activity was achieved with commonly prescribed drugs for patients with DS^39^, while drugs such as sodium channel blockers, increased seizure activity. This model is particularly advantageous as high throughput with short assay times is achieved, allowing rapid screening of novel compounds, a significant advantage over mouse models. In fact, lorcaserin, which has shown efficacy in some DS patients, was identified in zebrafish larvae^40^, further demonstrating this model has a role in the discovery of treatments for DS.

Overall, this study sets a foundation for screening novel compounds developed for DS through the use of the *Scn1a*^*A1783V/WT*^ model. While the *Scn1a*^*A1783V/WT*^ mouse did not respond to all drugs used as treatments for DS, the hyperthermia-induced seizure phenotype was highly pharmacoresistant, suggesting this model could lead to the discovery of novel and robust compounds for the treatment of DS, especially when paired with CLB and VPA treatment, as is done clinically.

## Acknowledgements

The authors thank the Epilepsy Therapy Screening Program at the National Institutes of Neurological Disease and Stroke for their review and comments on this manuscript. The authors would like to thank Jennifer Kearney for helpful discussions. This project has been funded by Federal funds from the National Institute of Neurological Disorders and Stroke, Epilepsy Therapy Screening Program, National Institutes of Health and Department of Health and Human Services, under Contract No. HHSN271201600048C.

