## Supporting Figure for "Development of an Antiseizure Drug Screening Platform for Dravet Syndrome at the NINDS contract site for the Epilepsy Therapy Screening Program"

**Supplemental Material**

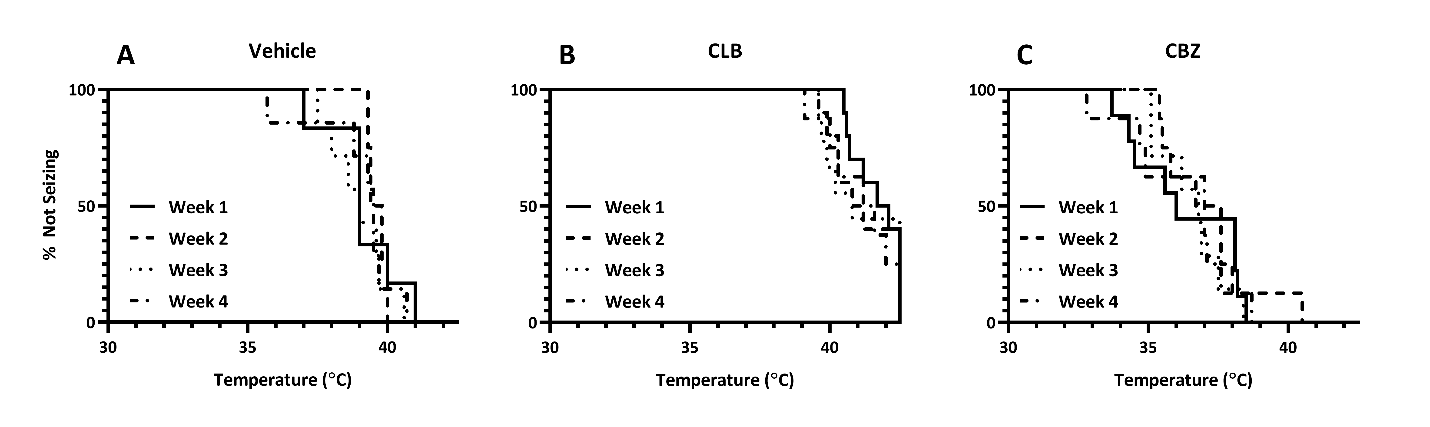

**Supporting Figure 1. Repeated treatment and hyperthermia induced seizures does not affect temperature threshold. A)** Vehicle treated mice were compared each week and there was no significant difference at temperatures at which Scn1a^A1783V/WT^ mice seized (Week 1 vs. Week 2: p = 0.8992, Week 1 vs. Week 3: p p = 0.4841, Week 1 vs. Week 4: p = 0.7405, Week 2 vs. Week 3: p = 0.5859, Week 2 vs. Week 4: p = 0.7812, Week 3 vs. Week 4: p = 0.6287, Week 1: n = 6, Week 2: n = 4, Week 3: n = 6, Week 4: n = 7, Log-rank (Mantel-Cox)). **B)** CLB treatment did not significantly affect temperature threshold when compared across treatment weeks (Week 1 vs. Week 2: p = 0.6385, Week 1 vs. Week 3: p = 0.7132, Week 1 vs. Week 4: p = 0.3770, Week 2 vs. Week 3: p = 0.9223, Week 2 vs. Week 4: p = 0.7327, Week 3 vs. Week 4: p = 0.7361, Week 1: n = 10, Week 2: n = 10, Week 3: n = 9, Week 4: n= 8, Log-rank (Mantel-Cox). **C)** CBZ treatment did not significantly alter temperature threshold when compared across treatment groups (Week 1 vs. Week 2: p = 0.8820, Week 1 vs. Week 3: p = 0.7798, Week 1 vs. Week 4: p = 0.9891, Week 2 vs. Week 3: p = 0.4373, Week 2 vs. Week 4: p = 0.4144, Week 3 vs. Week 4: p = 0.6950; Week 1: n = 9, Week 2: n =8, Week 3: n = 7, Week 4: n = 8, Log-rank (Mantel-Cox).

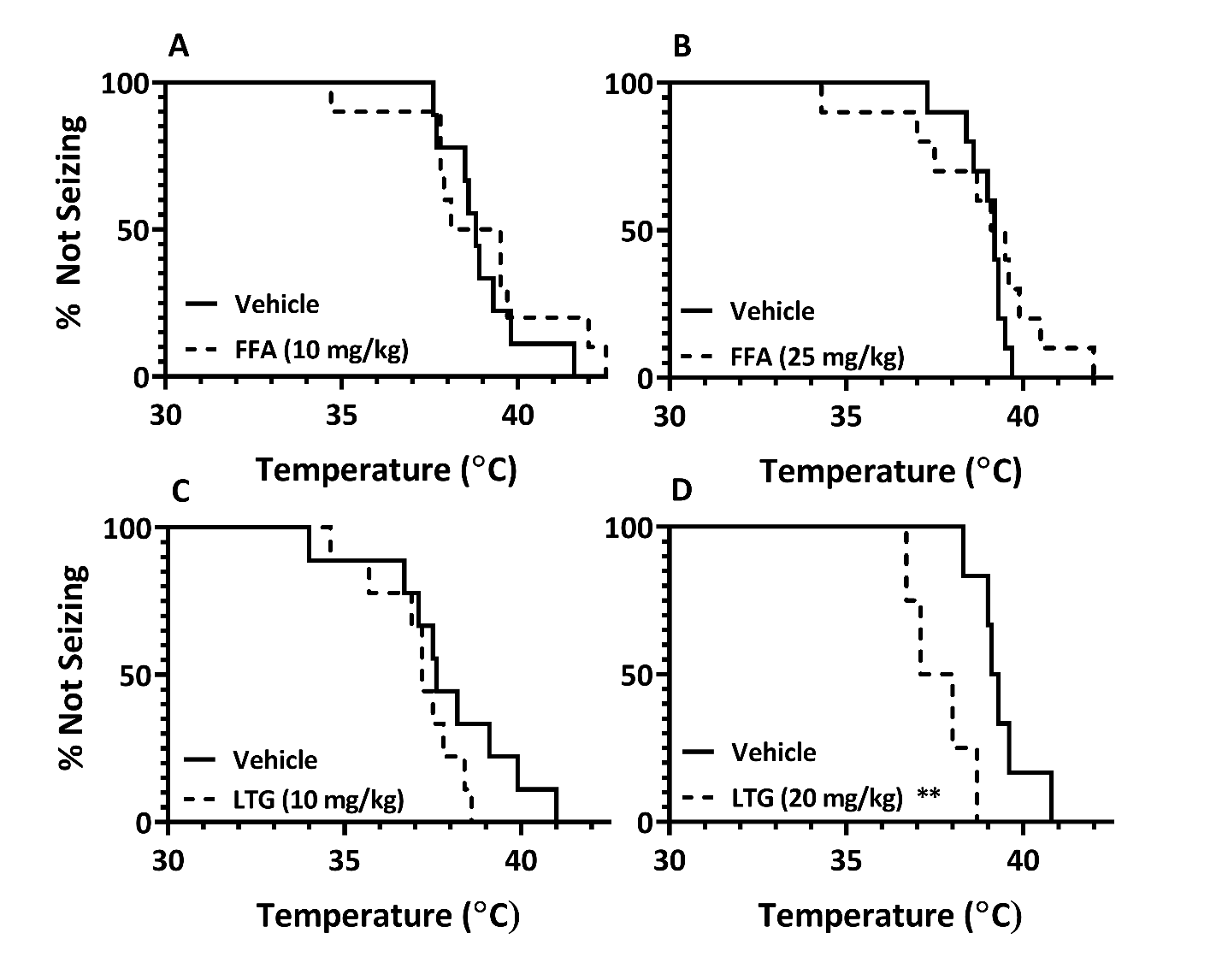

**Supporting Figure 2. Evaluation of FFA.** FFA was evaluated at 2 doses to determine the effect of each dose on hyperthermia-induced seizures. **A)** At a dose of 10 mg/kg, FFA did not significantly increase the temperature at which mice had hyperthermia-induced seizures (p=0.4862) as compared to vehicle treated mice. Log-rank (Mantel-Cox); Vehicle: n=10; FFA (10 mg/kg): n=10 **B)** 25 mg/kg FFA did not significantly affect the temperature threshold as compared to vehicle (p=0.2480) treated mice; Log-rank (Mantel-Cox); Vehicle: n=10; FFA (25 mg/kg): n=10. **C)** LTG (10 mg/kg) was administered as a single dose and had no effect on the temperature threshold (p=0.1898). Log-rank (Mantel-Cox); Vehicle: n=8, LTG (10mg/kg): n=9 **D)** LTG (20 mg/kg) was administered once a day over the course of 5 days and on Day 5, hyperthermia-induced seizure testing was conducted. Sub-chronic administration of LTG significantly lowered the temperature threshold at which mice seized (p=0.0046). Two mice did not undergo hyperthermia-induced seizure testing as they died as a result of LTG sub-chronic treatment. ** p<0.01, Log-rank (Mantel-Cox); Vehicle: n=6; LTG: n=4.
